# 25-Hydroxyl-cholesterol Binds and Enhance Anti-viral Effects of Zebrafish Monomeric C-reactive Proteins

**DOI:** 10.1101/371856

**Authors:** Melissa Bello-Perez, Alberto Falco, Jose Antonio Encinar, Beatriz Novoa, Luis Perez, Julio Coll

**Affiliations:** Instituto Nacional Investigaciones y Tecnologías Agrarias y Alimentarias, Dpto. Biotecnología. INIA. Madrid, Spain; Instituto de Biología Molecular y Celular, Universidad Miguel Hernández (IBMC-UMH). Elche, Spain; Investigaciones Marinas.CSIC. Vigo, Spain

**Keywords:** c-reactive protein, isoforms, 25-hydroxy-cholesterol, docking, zebrafish, SVCV

## Abstract

C-reactive proteins (CRP) are among the faster acute-phase inflammation-responses coded by one gene in humans (*hcrp*) and seven genes (*crp1-7*) in zebrafish (*Danio rerio*). In this study, preferential 25-hydroxycholesterol (25HOCh) binding to zebrafish CRP1-7 compared to other lipids were predicted by *in silico* docking and confirmed by solid-phase binding-assays. In addition, 25HOCh enhanced methyl-betacyclodextrin-sensitive (Cholesterol-dependent) CRP1-7 anti-viral effects in a fine-tunned isoform-dependent manner. *In silico* and structural studies suggested that the crosstalk between the anti-viral enhancements of both 25HOCh and CRP1-7 were dependent on protein monomers rather than oligomers but differred among isoforms. The presence of oxidized cholesterols in human atherosclerotic plaques amplifies the importance that similar interactions may have for vascular and/or neurodegenerative diseases during viral infections. In this context, the zebrafish model offers a genetic tool to further investigate how the expression and functions of different CRP isoforms and/or transcript variants may be controlled.

## Introduction

Previous studies have shown that in contrast to the one gene-coded human c-reactive protein (hCRP) [1], seven genes coded for CRP1-7 isoforms in zebrafish (*Danio rerio*) [2]. The hCRP are one of the most important clinical biomarkers for inflammation and most recently have been associated with relevant diseases such as those caused by cardiovascular and neurodegenerative disorders [3-5]. Circulating CRP molecules are planar oligomers of ~ 25 KDa monomers. The hCRP is pentameric (p-hCRP) while zebrafish CRP5 crystalizes as trimers [6]. It is not yet known whether or not other CRP1-7 isoforms are trimeric and what are their prevalent physiological conformation(s). On the other hand, some CRP1-7 isoform-dependent heterogeneous biological properties have been most recently described [6, 7].

The planar p-hCRP shows both lipid-recognition and functional-effector faces [5]. The recognition face binds phosphocholine heads exposed at the surface of prokaryotic/eukaryotic membranes in a Ca^++^ [8, 9] and phospholipase A_2_ [10] -dependent manner in damaged tissues [5]. Triggered by the CRP-Ca^++^-phosphocholine complexes, the functional-effector face binds C1q to activate the classical complement pathway, immunoglobulin Fc receptors to activate phagocytosis [11, 12] and other ligands to activate multiple cellular functions [10]. In order to accomplish all that variety of functions, hCRP shows, at least, 4 different conformations [5, 13]: **i)** Inactive serum-circulating p-hCRP, which is present in low concentrations in healthy humans, increasing 100-1000-fold after inflammation, **ii)** Pro-inflammatory tissue-associated p-hCRP* [4], **iii)** Pro-inflammatory tissue-associated monomeric hCRP (m-hCRP) with wider ligand capacities which include cholesterol (Ch) [14-16] and **iv)** Disulphide-reduced m-hCRP which activates lymphoid and many other cells [5, 16-18]. Despite different oligomeric structures between p-hCRP and t-CRP5 [6], their protein hydrophobic profiles, two cysteine residues per monomer, Ca^++^-binding amino acids and location of phosphocholine (PC)-binding pockets, were highly conserved [19]. On the other hand, previous transcriptomic studies on *crp1-7* genes demonstrated differential transcript expression in zebrafish tissues [2], in survivors of viral infection [20] and in mutants defective in adaptive immunity [21]. In addition, unexpected *crp1-7*/CRP1-7 isoform-dependent anti-viral *in vitro* and *in vivo* activities have been described showing also that *crp2/*CRP2 and *crp5/*CRP5 transcripts/proteins were the most modulated during *in vivo* viral infections in contrast to *crp1/*CRP1/ and *crp7*/CRP7. These recent findings unrevealed novel anti-viral CRP1-7 direct or indirect activities in zebrafish which, to our knowledge, have not been described yet for hCRP. All the similar properties mentioned above suggested analogous biological functions for p-hCRP and CRP1-7 [7], however, whether CRP1-7 isoforms do physiologically exist as different oligomeric structures, conformations and/or become specialized in different ligand-binding or biological functions remains unexplored.

In this work, we focused by *in silico* and *in vitro* studies on the lipid-docking/binding, anti-viral activities and oligomeric forms of the CRP1-7 isoforms and CRP5 transcript variants of zebrafish. We have found that **i)** Ca^++^-independent docking/binding of CRP1-7 to Ch was preferred to other lipids, **ii)** HOChs were a preferential target for CRP1-7, specially for CRP1, **iii)** HOChs enhanced the anti-viral direct or indirect effects by zebrafish CRP1-7 in an isoform-dependent manner and **iv)** CRP2 / CRP5 and numerous CRP5 transcript variants have an stronger tendency to fold as trimers. The abundance of oxidized Chs in atherosclerotic plaques amplifies the importance that similar interactions may have for vascular diseases and neurodegenerative disorders during viral infections in humans [22-24]. The functional significance of the CRP oligomer-monomer conversion (and *viceversa*?) need to be further clarified to evaluate new chemotherapeutic targets [10, 25]. Zebrafish may offer a good genetic model to continue such studies.

## Material and Methods

### *In silico* docking predictions between zebrafish CRP1-7 and lipids

The AutoDock Vina [26] included in the PyRx program package [27] was used to predict Gibbs free-energy of docking (ΔG) of 60 x 60 x 60 Å grids surrounding the CRP1-7 molecules. When required for comparison purposes with experimental data, the output ΔG energies were converted to constant inhibition (Ki) in molar concentrations (M), by the formula Ki = exp([ΔG x 1000] / [R x T]) (R = 1.98 cal/mol, and T = 298 °C)[28]. The predicted structures were visualized in PyRx and/or PyMOL (https://www.pymol.org/).

### Cell culture in EPC cell monolayers

The *epithelioma papulosum cyprinid* (EPC) cells from fathead minnow fish (*Pimephales promelas*) were obtained from the American Type Culture Collection (ATCC, Manassas, Vi, USA, code number CRL-2872). EPC cell monolayers were grown at 26 °C in a 5 % CO_2_ atmosphere in RPMI-1640 Dutch modified cell culture medium (Gibco, UK) supplemented with 20 mM HEPES, 10 % fetal bovine serum, FBS (Sigma, St. Louis, USA), 1 mM piruvate, 2 mM glutamine, 50 μg/ml of gentamicin (Gibco) and 2 μg/ml of fungizone [21].

### Preparation of spring viremia carp virus (SVCV)

The isolate 56/70 of Spring Viremia Carp Virus (SVCV) from carp *C. carpio* [29], recently renamed *Carp Sprivivirus* [30], was replicated in EPC cell monolayers at 26 °C, in the cell culture media described above except for 2 % FBS and absence of the CO_2_ atmosphere. Supernatants from SVCV-infected EPC cell monolayers were clarified by centrifugation at 4000 g for 30 min and kept at −80 °C until used [21].

### Estimation of the effects of methyl-betaciclodextrin (MBCD) in SVCV infectivity

EPC cell monolayers treated for 2 h with different concentrations (0-8 mM) of MBCD were incubated with SVCV for 24 h and assayed for fluorescent focus forming units (ffu) (see later). The results were expressed as percentage of SVCV infectivity calculated by the formula, 100 x (ffu treated with MBCD / ffu non-treated with MBCD). To assay for viability, EPC cell monolayers treated with different concentrations of MBCD as above were incubated with 0.5 mg/ml of 3-(4,5-dimethylthiazol-2-yl)-2,5-diphenyltetrazolium bromide (MTT) in a Krebs–Hensleit–HEPES buffer (115 mM NaCl, 5 mM KCl, 1 mM KH_2_PO_4_, 1.2 mM MgSO_4_, 2 mM CaCl_2_, and 25 mM HEPES at pH 7.4) for 3 h, absorbance at 570 nm measured and the percentage of viability calculated by the formula, absorbance of MBCD treated cells / absorbance of untreated cells. Means and standard deviations (n=2) were interpolated and smoothed by the cubic B-spline method in Origin Pro 2017 (Northampton, MA, USA).

### Construction of recombinant pRSET-CRP1-7 for *E.coli* expression

The corresponding mRNA sequences of the CRP1-7 proteins [7] were used for the design, construction and expression in *E.coli*. All the corresponding synthetic DNA sequences were cloned into the pRSET adding poly-histidine tails (polyH) at their C-terminal ends (GeneArt, Regensburg, Germany). The purified plasmids were then transfected into *E.coli* BL21(DE3) and grown at 37 °C. The resulting recombinant bacteria from the pRSET-*crp1-7* constructs were induced with IPTG at 25 °C. Gradient 4-20 % polyacrylamide gel electrophoresis (PAGE) and Western blot were used to detect CRP1-7 expression.

### Construction of recombinant rCRP1, rCRP2, rCRP5, rCRP7 for insect expression

The mRNA sequences of the CRP1, CRP2, CRP5 and CRP7 described before [7] were used for the design, construction and expression in insect cells (GenScript, Piscataway, NJ, USA). Briefly, target DNA containing the gp67 signal peptide + CRP1-7 + Flag (DYKDDDK) + 6 x polyHis sequences (construct size of ~3 Kbp) were synthesized and sub-cloned into the pFastBac1^TM^ baculovirus transfer vector (Invitrogen). The pFastBac1 recombinants were transfected into DH10 Bac^TM^-competent *E. coli* cells and bacmids prepared from selected *E. coli* clones. Next, recombinant baculoviruses were generated in *Spodoptera frugiperda* (Sf9) insect cells. For that, Sf9 cells cultured in Grace’s insect media (Gibco BRL) with 10 % foetal bovine serum, 3 % non-essential amino acids and 20 μg/ml gentamicin at 28 °C [31] were co-transfected with bacmids and baculovirus using Cellfectin II. Supernatants containing the recombinant baculoviruses were obtained 72 h post-transfection with titres of ~ 10^7^ pfu/ml.

For rCRP expression and purification, 500 ml of Sf9 cell supernatants were harvested 72 h post-infection and dialyzed against 50 mM Tris, pH 8.0, 500 mM NaCl. The rCRP-containing medium were incubated with Flag or Ni^++^ columns equilibrated with 50 mM Tris, 500 mM NaCl, 5 % glycerol, pH 8.0, eluted with 200 μg/ml of the Flag peptide or 150 mM imidazole, dialyzed against equilibration buffer and kept at −20 °C until used. Purified rCRP were loaded in 8-20 % SDS-polyacrylamide gels (BioRad), electrophoresed, and transferred to nitrocellulose membranes (Schleicher & Schuell) to detect specific tag epitopes. The membranes were blocked with phosphate buffer saline-0.05 % Tween 20 with 4 % skim milk, incubated with anti-poly-H monoclonal antibody MAb (Sigma) for 1 h, and then with anti-mouse horseradish peroxidase-conjugated immunoglobulins (Sigma) and visualized with diaminobenzidine (DAB). Protein concentrations were determined using the bicinchoninic acid (BCA) method [32] and confirmed by PAGE with BSA as standard.

### Production of rabbit antibodies to recognize zebrafish CRP1-7 isoforms

To detect CRP1-7 isoforms in lipid-binding assays and after PAGE by Western blotting, anti-CRP1-7 rabbit antibodies were raised (GenScript, Piscataway, NJ, USA) against 3 of the longest more conserved amino acid stretches such as peptide p1 (^18^SYVKLSPEKPLSLSAFTLC), peptide p2 (^189^DWDTIEYDVTGN) and peptide p3 (^129^RPGGTVLLGQDPDSYVGSFC). All p1, p2 and p3 were located at the CRP1-7 surface, as shown by PyMOL modelling of trimeric CRP5 ± Ca^++^ (4PBP.pdb and 4PBO.pdb, respectively) [6] (not shown). To reduce assay backgrounds, the anti-peptide antibodies were purified by affinity chromatography against the corresponding synthetic peptides coupled to CNBr-activated Sepharose. Only the affinity-purified anti-p3 antibodies bound purified insect-made rCRP2, rCRP5 and rCRP7 on Western blots under denaturing and non-denaturing conditions (Figure 1A) and recognized EPC cells transfected with pMCV1.4-*crp2-7* by immunofluorescence (not shown).

**Figure 1.**
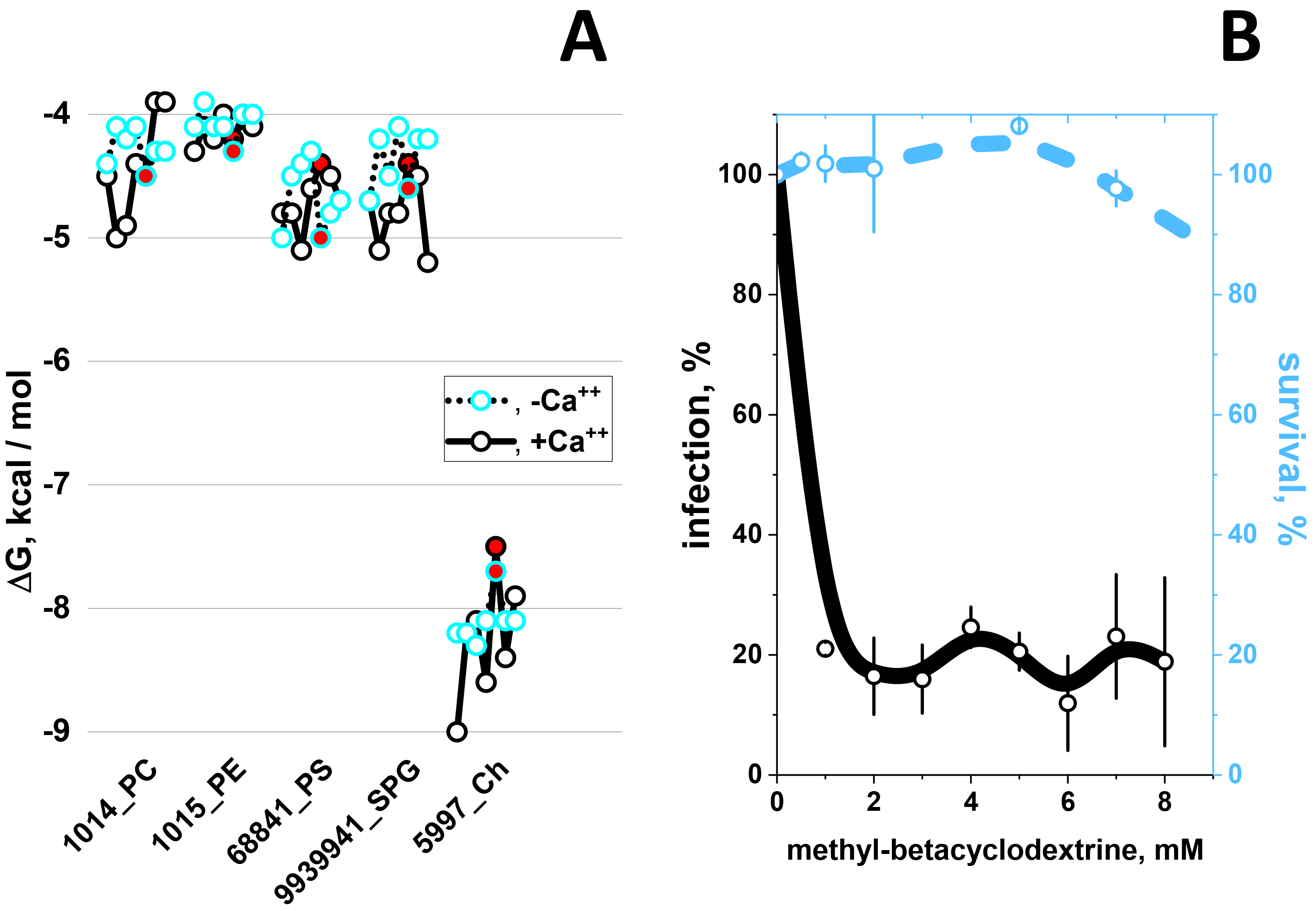
CRP1-7 preferential docking to Ch (A) and inhibition of SVCV infectivity by methyl-betacyclodextrin (MBCD) (B) **A)** Docking predictions to selected lipid-heads and Ch. The CRP1-7 were SWISS-modelled using as templates the CRP5 (GenBank accession number JF772178.1), 4PBP.pdb (**+Ca^++^**) and 4PBO.pdb (**-Ca^++^**) 3D-structures [6]. Structures were extracted from *.sdf from PubChem (https://pubchem.ncbi.nlm.nih.gov/search/search.cgi) and converted to *.pdbqt using the Babel program from the PyRx package [27]. ΔG values from two independent dockings were averaged, the standard deviations (<5%) omitted for clarity. **PC**, Phosphocholine. **PE**, Phosphoethanolamine. **PS**, Phosphoserine. **SPG**, palmitoyl sphingomyelin. **Ch,** cholesterol. **Numbers before the names**_, PubMed ID numbers. **Blue open circles**, consecutive CRP1-7 isoforms from left to right in the absence of Ca^++^. **Black open circles**, consecutive CRP1-7 isoforms from left to right in the presence of Ca^++^. **Red circles**, CRP5. **Black lines**, +Ca^++^. **Dot lines**, -Ca^++^. **B)** Effect of methyl-betacyclodextrin (MBCD) on SVCV infectivity. MBCD-treated EPC cell monolayers were incubated with SVCV for 24 h and assayed by ffu. The results were expressed as infectivity percentages calculated by the formula, 100 x (ffu treated with MBCD / ffu non-treated with MBCD). To assay for viability, MBCD-treated EPC cell monolayers were incubated with MTT for 3 h, absorbance at 570 nm measured and the percentage of viability calculated by the formula, absorbance of treated cells / absorbance of untreated cells. **Open blue or black circles and their vertical lines**, means and standard deviations (n=2), respectively. The data were then interpolated and smoothed by the cubic B-spline method in Origin Pro 2017 (Northampton, MA, USA). **Black line,** SVCV infectivity. **Blue dashed line**, EPC cell monolayer viability.

### Binding of CRP1-7 to solid-phase lipids

Binding of CRP1-7 to lipids were assayed in solid-phase 96-well plates (Nunc, Maxisorb) by modifying previously described methods [33]. Wells were coated to dryness with several concentrations of ethanol dissolved lipids and kept dried until used. To assay for CRP1-7 binding, the plates were first washed with 0.1 M sodium borate, 1 mM Ca_2_Cl buffer, pH 8, and then 0.5 μg/well of rCRPs or 10-fold diluted ssCRP1-7 added in 50 μl of the same buffer and incubated for 60 min. After washing, bound CRP1-7 were detected with rabbit anti-p3 and peroxidase-labelled goat anti-rabbit IgG. Peroxidase was finally assayed with OPD as described before [34, 35]. All the resulting data were interpolated and smoothed by the cubic B-spline method in Origin Pro 2017 (Northampton, MA, USA).

### Binding of CRP5 pepscan peptides to solid-phase 25HOCh and docking predictions

A series of 15-mer peptides overlapping 5 amino acids of the CRP5 sequence were chemically synthesized by adding an amino-terminal biotin molecule (Chiron Mimotopes, Victoria, Australia). The synthetic pepscan peptides were diluted in distilled water to 4 mg/ml and kept frozen until use.

To perform the binding experiments, 2 μg of 25HOCh were dissolved in 50 μl of ethanol and dried into polystyrene wells of 96-well Nunc Maxisorb plates. After washing the plates with 0.1 M borate buffer pH 8, 1 mM CaCl_2_, pepscan peptides (0.05 μg in 50 μl) were added to each of the wells and incubated for 60 min. After washing, 1000-fold diluted peroxidase-labelled streptavidin were added and incubated for 30 min. After the last washing, OPD was used to detect the amount of peroxidase as described before [35].

To perform the *in silico* docking predictions, the best modelled CRP5 pepscan peptide sequences predicted in solution by the Mobyle program http://mobyle.rpbs.univ-paris-diderot.fr/cgi-bin/portal.py#forms::PEP-FOLD [36] were docked to all possible predicted conformations of 25HOCh. All the resulting docking data were interpolated and smoothed by the cubic B-spline method in Origin Pro 2017 (Northampton, MA, USA) and the data which best fitted pepscan binding selected for representation. Validation of such strategy was confirmed by the high correlation obtained among similarly modelled VHSV G protein pepscan 15-mer peptides and previously published binding data to labelled phosphatidylserine [37] and phosphatidylinositol-bisphosphate [38] (not shown).

### Preparation of pMCV1.4 plasmids coding for *crp1-7*

Each of the chemically synthesized *crp1-7* and green fluorescent protein (*gfp*) genes was subcloned into the pMCV1.4 plasmid as described before [7]. The resulting pMCV1.4-*crp1-7* and pMCV1.4-*gfp* plasmid constructs were used to transform *E.coli* DH5alpha, amplified and isolated with the Endofree Plasmid Midi purification Kit (Qiagen, Germany) according to manufacturer’s instructions. Purified plasmid solutions containing 80-100 % of plasmid DNA, as shown by agarose gel electrophoresis were stored at −20 °C.

### Preparation of CRP1-7 enriched supernatants

To produce ml amounts of CRP1-7 enriched supernatants (ssCRP1-7), 60 % confluent EPC cell monolayers in 25 cm^2^ bottles in 5 ml of cell culture medium were transfected with 5 μg of each of the pMCV1.4-*crp1-7* plasmids complexed with 15 μl of FuGENE HD (Promega, Madison, WI, USA) for 24 h at 22 ° C (transfection efficiency of 15.2-30.4 %, n=3 as estimated by transfection with pMCV1.4-*gfp*). After washing with fresh cell culture medium, the ssCRP1-7 were harvested 3-days later, cell debris eliminated by centrifugation, sterilized by filtration through 0.2 μ filters and kept in aliquots at −80 °C until used, essentially following the previously described procedure [7].

### SVCV infection of pre-incubated EPC cell monolayers with 25HOCh and ssCRP1-7

To detect the effects of 25HOCh and ssCRP1-7 (25HOCh + ssCRP1-7) on SVCV infection, the concentrations of 25HOCh, and ssCRP1-7 and the multiplicity of infection (m.o.i.) of SVCV were first optimized (not shown). Optimal conditions were obtained when the EPC cell monolayers were pre-incubated with 100 μl of 4-fold diluted ssCRP1-7 or ssGFP in RPMI with 2% FBS in the absence or presence of 10 μM 25HOCh for 20 h at 26 °C, the monolayers washed twice and SVCV added at 10-2 m.o.i. To estimate the extent of SVCV infection, the monolayers were incubated with SVCV for 2 h, washed, and incubated for 24 h at 26 °C. The number of infected EPC cells was estimated by flow cytometry after staining with monoclonal anti-SVCV (BioX Diagnostics SA, Jemelle, Belgium) and fluorescein-labelled goat anti-mouse immunoglobulins as described before [7]. The number of EPC infected cells varied from 29.6-39.7 % or 9.9-20.1 % (n=3) after pre-incubation of the EPC cell monolayers with either 25HOCh or ssCRP1-7 alone, respectively. The results of pre-incubation with 25HOCh + ssCRP1-7 were expressed in relative percentage of infection ± 25HOCh as calculated by the formula, 100 x (percentage of infected EPC cells pre-incubated with 25HOCh + CRP1-7 / percentage of infected EPC cells pre-incubated in absence of 25HOCh and presence of CRP1-7).

### *In silico* modelling of CRP1-7 tridimensional structures

To explore the CRP1-7 tridimensional structures, their protein sequences were automatically modelled (RMSD<0.3 Ă) using the SWISS-MODEL homology server (https://swissmodel.expasy.org/interactive) [39-41]. Tridimensional structures of the target CRP1-7 sequences were predicted after pairwise comparing interfaces between the target and the best template selected by the program. For each possible interface with > 10 residue-residue interactions, the QscoreOligomer score was calculated and averaged from all predicted interfaces [39, 42]. The templates that resulted selected for the automatical modelling corresponded to zebrafish CRP5 ± Ca++ (4PBP.pdb and 4PBO.pdb) [6] (RCSB data bank at http://www.rcsb.org/pdb/home/home.do).

## Results and discussion

### Preferential docking predictions of zebrafish CRP1-7 to Ch

To predict their docking ΔG energies to CRP1-7, phosphocholine head (PC), other phospholipid heads [43-45] and cholesterol (Ch) [16] were selected because of their hCRP ligand properties. Interestingly, the results predicted the lowest ΔG (prediction of stronger binding) for Ch (ΔG ranges from −7.5 to −9 Kcal/mol) compared to phospholipid-heads (ΔG ranges from −4 to −5.5 Kcal/mol). The addition of a glycerol molecule to those phospholipid-heads did not changed their predicted ΔG (not shown). Results also predicted Ch docking energies more Ca^++^-independent than most other lipid-heads (Figure 1A), and alternative docking locations for Ch and other lipid-heads (not shown).

### Membrane Ch sequestering by methyl-betacyclodextrins decreases SVCV infection

To explore whether Ch was implicated in any anti-viral effects, SVCV infections were studied after previous treatment of the EPC cell monolayers with methyl-betacyclodextrins (MBCD), a sequestering agent for membrane Ch. Treatment with MBCD from 0.5 to 8 mM lowered the SVCV infectivity to ~ 20 % (Figure 1B, black line), while those concentrations did not have any significative effects on cell survival (Figure 1B, blue dashed line). These results confirmed that the Ch presence in the cell membranes was required for SVCV infectivity and suggest that the CRP1-7-Ch predicted docking may have anti-viral effects. Because, **i)** Ch-containing lipid rafts participate in interactions with hCRP [46], **ii)** Ch is a key molecule involved in coronary diseases and **iii)** Ch-related physiological compounds are highly diverse, an screening for other physiological Ch-related compounds was performed before studying any possible anti-viral effects.

### Preferential docking predictions of zebrafish rCRPs to hydroxycholesterols (HOChs)

When 23 Ch-related physiological compounds were explored for docking to CRP1-7, the stronger predictions (ΔG ranges between −7.5 to −9.3 Kcal/mol) were obtained for most hydroxy derivatives for most CRP1-7 (Figure 2). The majority of the lowest ΔG values were obtained for CRP1 while CRP5 showed ~ 0.5-1 Kcal/mol higher ΔG than CRP1, depending on the Ch-related molecule. The most relevant results of these Ch-related docking predictions could be summarized as follows: **i)** Water-soluble hydroxy Ch derivatives (HOChs) interacted with CRP1-7 within lower ΔG ranges (−8.0 to −9 Kcal/mol), **ii)** Most of the lower ΔG values corresponded to CRP1, while most of the highest ΔG corresponded to CRP5, and **iii)** The 25-hydroxycholesterol (25HOCh) was unique among the tested HOChs since docking predicted the lowest ΔG values (~ −9 Kcal/mol).

**Figure 2.**
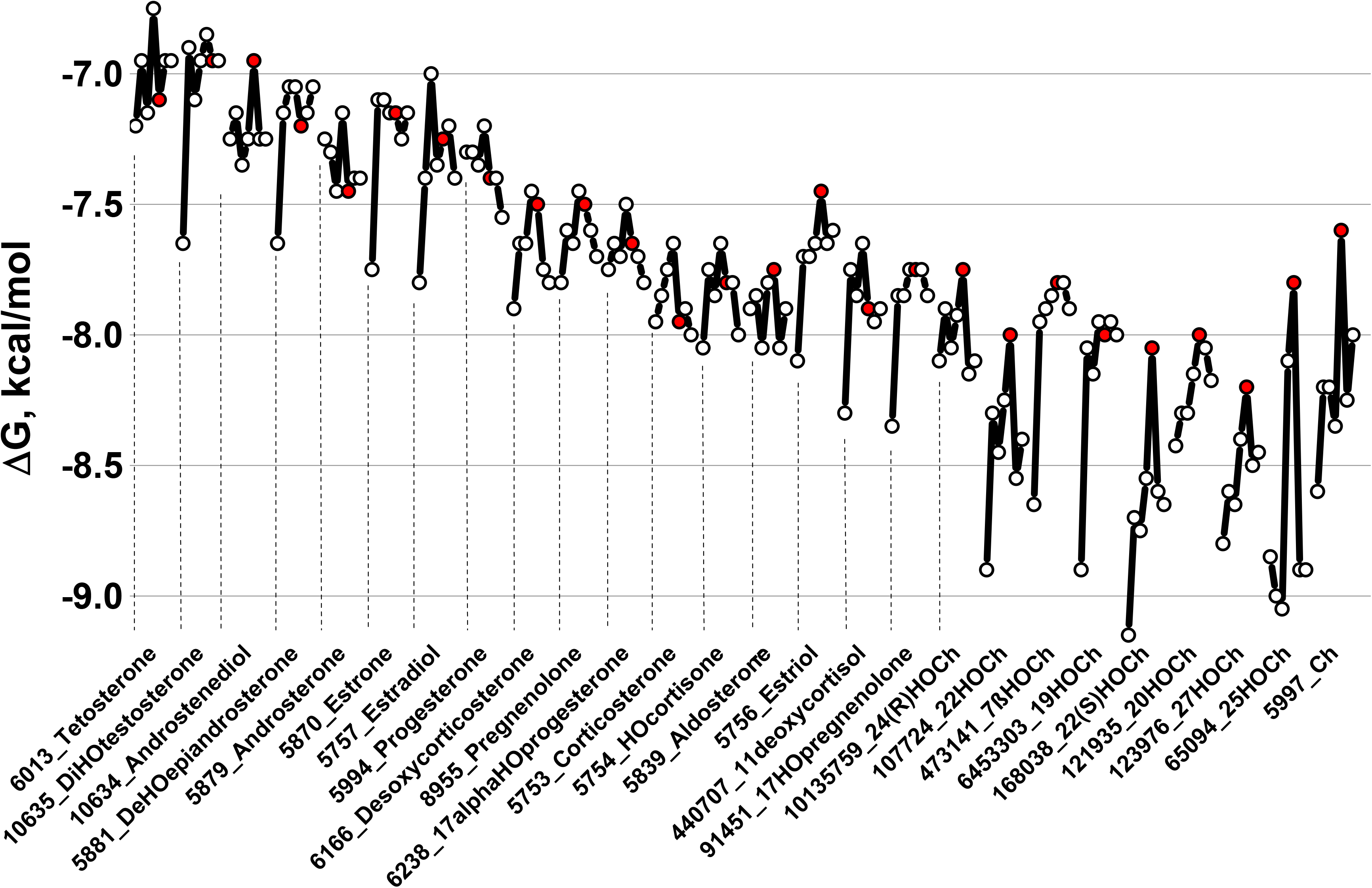
CRP1-7 docking predictions to several Ch-related physiological molecules. CRP1-7 models, Ch-related physiological molecules and ΔG predictions were obtained as described in the legend of Figure 3. Because the predicted ΔG values in the absence or presence of Ca^++^ were similar, only the mean ΔGs ± Ca^++^ were represented. **Open circles**, consecutive CRP1-7 isoforms from left to right. **Red circles**, CRP5. **Numbers before the names**_, PubMed ID numbers. **HO**, hydroxy. **Ch,** cholesterol.

To explore for the existence of other possible Ch-related compounds with still lower ΔG values which could be used as anti-inflammatory chemotherapeutic drugs, a library of 1093 Ch-related synthetic molecules were docked to CRP1-7. The frequency distribution of predicted ΔGs showed the existence of Ch-related non-physiological compounds with lowest ΔG values from −13.3 to −12 Kcal/mol (mean - 3 x standard deviations = 12 Kcal/mol) (Table 1). Most of the new molecules identified contained Deuterium, Fluorine, Bromine or Chlorine atoms and 66.6 % contained, at least, one hydroxy group per molecule. Therefore, some of these newly identified Ch-related derivatives could be further used for drug applications and/or mechanistic studies in the future.

**Table 1.**
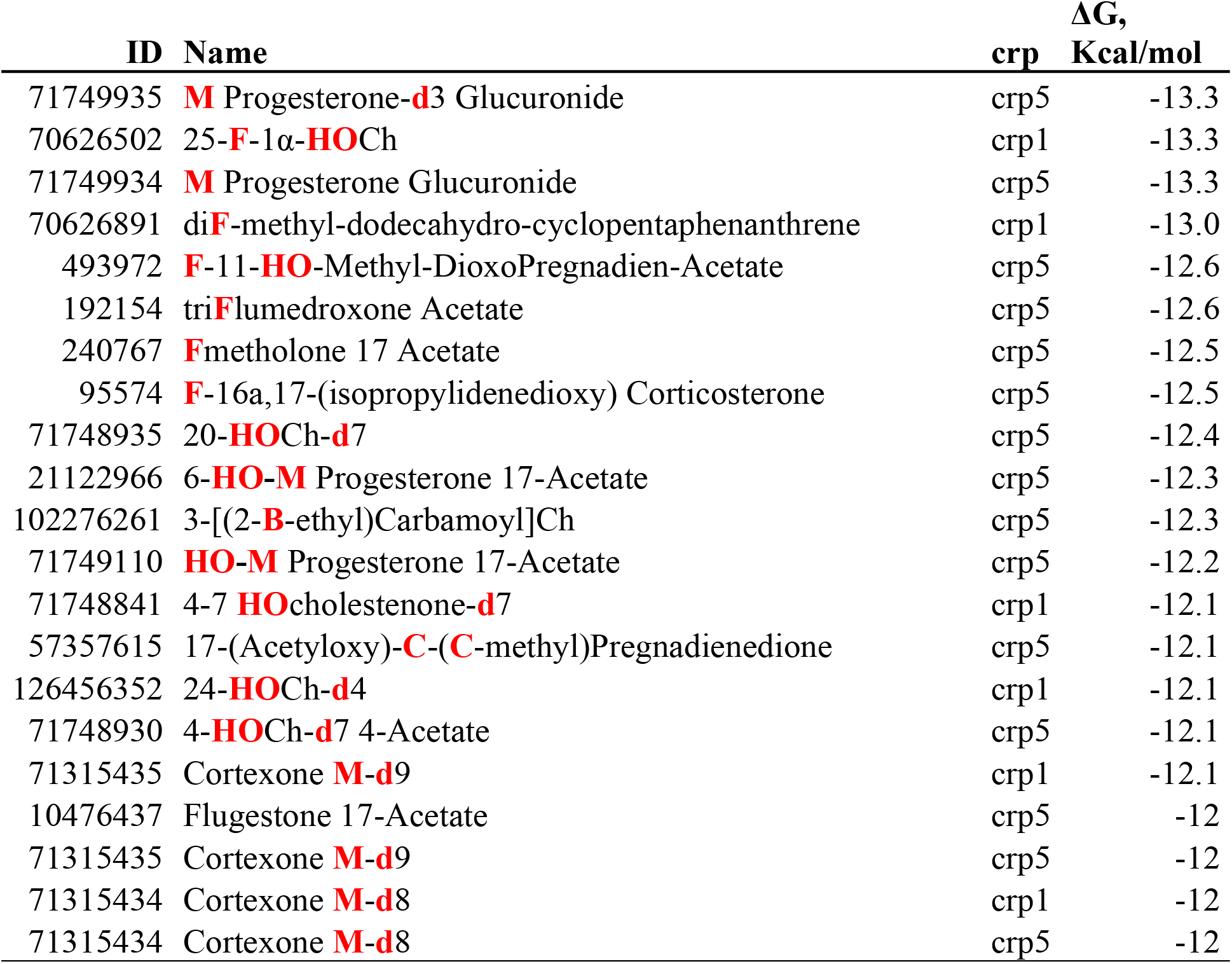
Ch-related non-physiological compounds with best docking predictions to CRP1-7. Ch-related non-physiological compound structures were retrieved from several libraries obtained from PubChem in a *.sdf format. To construct the library, 550 Chs, 314 Colestens, 73 Corticosterones, 41 Dehydroepiandrosterones (DHEAs), 107 Estriols, 99 Pregnenolones, 196 Progesterones and 107 HOChs were retrieved. Duplicated and extremely long molecules were eliminated from the total of 1487 *.sdf, resulting in a downsized library of 1093 *.pdbqt archives. After docking, the frequency distribution of ΔG showed two peaks with means at −11 and −7 Kcal/mol, respectively. Only those Ch-related compounds with ΔG < −12 Kcal/mol (mean + 3 x standard deviations) were tabulated. **ID,** PubMed number**. Red bold HO,** Hydroxy**. Red bold d,** deuterium**. Red bold F,** fluoro-**. Red bold C,** chloro-**. Red bold B,** Bromo. **Red bold M**, 17-acetyl-6,10,13-trimethyl-3-oxo-1,2,6,7,8,9,11,12,14,15,16,17-dodeca **HO**cyclopenta[a]phenanthren-16-yl) acetate (Medroxy).

| ID | Name | crp | $\Delta G$,<br>Kcal/mol |
| --- | --- | --- | --- |
| 71749935 | **M** Progesterone-**d3** Glucuronide | crp5 | -13.3 |
| 70626502 | 25-**F**-1α-**HO**Ch | crp1 | -13.3 |
| 71749934 | **M** Progesterone Glucuronide | crp5 | -13.3 |
| 70626891 | di**F**-methyl-dodecahydro-cyclopentaphenanthrene | crp1 | -13.0 |
| 493972 | **F**-11-**HO**-Methyl-DioxoPregnadien-Acetate | crp5 | -12.6 |
| 192154 | tri**F**lumedroxone Acetate | crp5 | -12.6 |
| 240767 | **F**metholone 17 Acetate | crp5 | -12.5 |
| 95574 | **F**-16a,17-(isopropylidenedioxy) Corticosterone | crp5 | -12.5 |
| 71748935 | 20-**HO**Ch-**d7** | crp5 | -12.4 |
| 21122966 | 6-**HO**-**M** Progesterone 17-Acetate | crp5 | -12.3 |
| 102276261 | 3-[(2-**B**-ethyl)Carbamoyl]Ch | crp5 | -12.3 |
| 71749110 | **HO**-**M** Progesterone 17-Acetate | crp5 | -12.2 |
| 71748841 | 4-7 **HO**cholestenone-**d7** | crp1 | -12.1 |
| 57357615 | 17-(Acetyloxy)-**C**-(**C**-methyl)Pregnadienedione | crp5 | -12.1 |
| 126456352 | 24-**HO**Ch-**d4** | crp1 | -12.1 |
| 71748930 | 4-**HO**Ch-**d7** 4-Acetate | crp5 | -12.1 |
| 71315435 | Cortexone **M**-**d9** | crp1 | -12.1 |
| 10476437 | Flugestone 17-Acetate | crp5 | -12 |
| 71315435 | Cortexone **M**-**d9** | crp5 | -12 |
| 71315434 | Cortexone **M**-**d8** | crp1 | -12 |
| 71315434 | Cortexone **M**-**d8** | crp5 | -12 |

### Binding of zebrafish rCRPs to hydroxycholesterols (HOChs), Ch and PC

We then tried to confirm some of the docking predictions by solid-phase binding assays. Because 25HOCh demonstrated the highest docking to CRP1-7 and because of its recently described anti-viral activities [24, 47], 25HOCh was compared to Ch / PC bindings (the former because containing the traditional ligand for hCRP). Using 25HOCh coated solid-phase polystyrene plates [33], the binding results confirmed the higher docking of rCRP5 / rCRP7 to Ch / 25HOCh compared to PC (Figure 3A). Binding of rCRP7 to Ch / 25HOCh were slightly higher than to rCRP2 or rCRP5 (Figure 3, rCRP7) while rCRP2 / rCRP5 binding to Ch or PC were relatively low (Figure 3, rCRP2 and rCRP5). To complete the study, we explored the rest of isoforms for binding to 25HOCh using CRP1-7-enriched supernatants from EPC cells transfected with pMCV.4-*crp1-7* (ssCRP1-7) as a source of CRP1-7. The results of these experiments showed different concentration-dependent profiles for different CRP1-7, being the CRP1 the most active at the lower 25HOCh concentrations assayed (<10 μM) (Figure 3B). On the other hand, although CRP7 showed slightly higher bindings at >100 μM 25HOCh, similar values were obtained for all ssCRP1-7 at those higher concentrations. Binding of ssCRP1-7 to solid-phase 25HOCh showed relatively lower values than to rCRPs, most probably due to the lower CRP concentrations in the ssCRP1-7 (compare ordinate values of Figure 3A and B).

**Figure 3.**
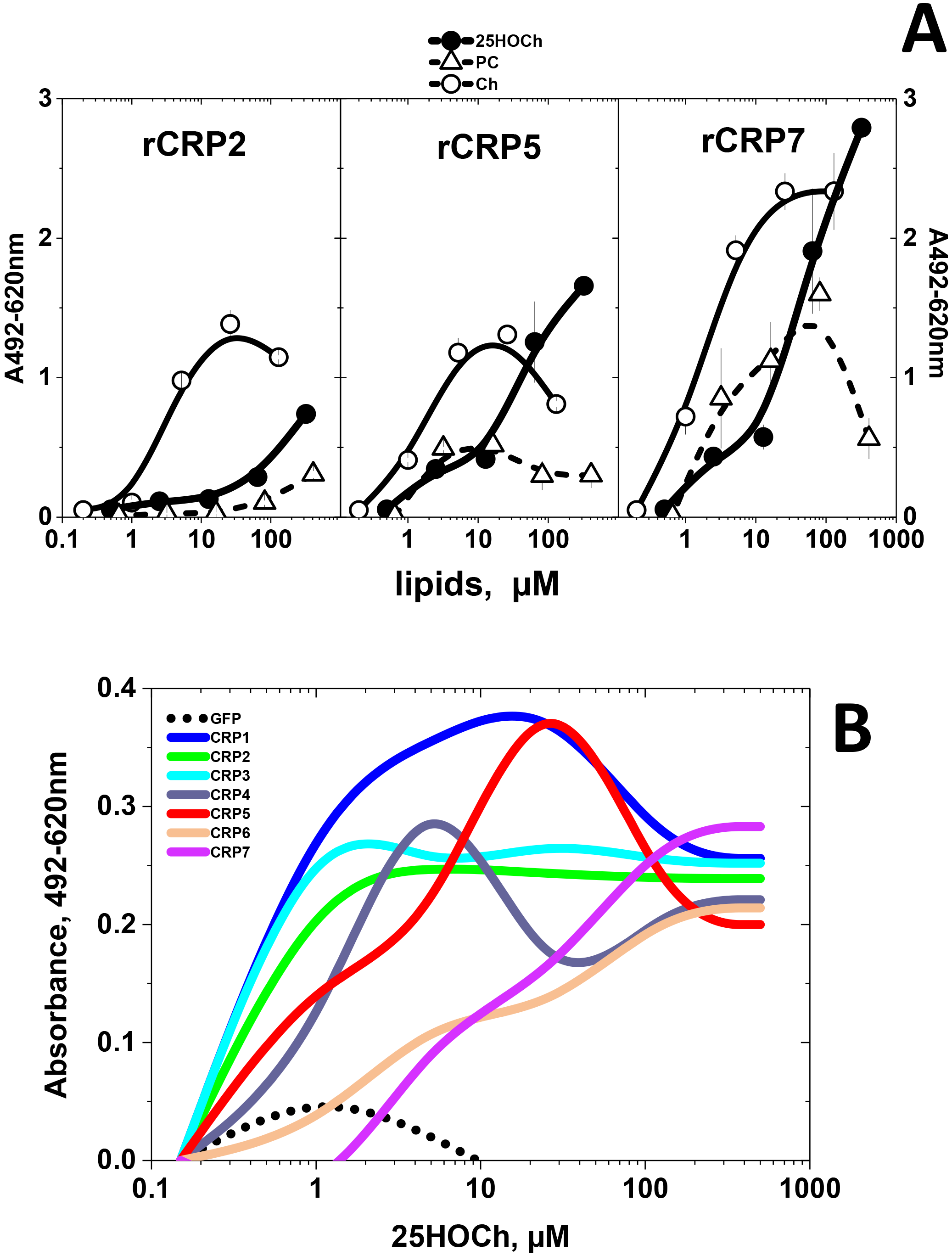
rCRP (A) and ssCRP1-7 (B) binding to solid-phase lipids. Binding of purified rCRPs and ssCRP1-7 to selected lipids were assayed by using solid-phase plates of 96-well (Nunc, Maxisorb) coated to dryness with several lipid concentrations dissolved in ethanol. The lipid-coated plates were washed with 0.1 M sodium borate, 1 mM Ca_2_Cl pH 8 (borate buffer) and incubated with rCRP2 or ssCRP1-7 in borate buffer for 1 h in 50 μl volume. To detect bound rCRP or ssCRP1-7, rabbit anti-CRP p3 peptide, peroxidase labeled goat anti-rabbit IgG and OPD were used as described before [34, 35]. Means and standard deviations from 2 independent experiments (n=2) were represented. **A)** rCRP at 0.5 μg/well in borate buffer. **Open triangles**, solid-phase phosphatidylcholine (PC). **Open circles**, solid-phase Ch. **Black circles**, solid-phase 25HOCh. **B)** ssCRP1-7 were 10-fold diluted in borate buffer. Means (n=6) were interpolated and smoothed by the cubic B-spline method in Origin Pro 2017 (Northampton, MA, USA). **Black points**, supernatant from pMCV1.4-*gfp* transfected cells. **Blue line**, supernatant from pMCV1.4-*crp1* transfected cells. **Green line**, supernatant from pMCV1.4-*crp2* transfected cells. **Light-blue line**, supernatant from pMCV1.4-*crp3* transfected cells. **Gray line**, supernatant from pMCV1.4-*crp4* transfected cells. **Red line,** supernatant from pMCV1.4-*crp5* transfected cells. **Orange line**, supernatant from pMCV1.4-*crp6* transfected cells. **Purple line**, supernatant from pMCV1.4-*crp7* transfected cells.

### Mapping of CRP5 binding and docking to 25HOCh

To further clarify the 25HOCh binding to CRP1-7, we mapped such interaction. Because m-hCRP rather than p-hCRP is the conformation that preferentially binds Ch [16, 17, 48], some non-conformational motifs may conserve Ch- / 25HOCh-binding activity. Therefore, we selected a pepscan to explore for possible non-conformational interactions with 25HOCh by solid-phase binding assays and by docking predictions.

For the peptide binding assays, each of the synthetically biotinylated 15-mer peptides derived from the CRP5 amino acid sequence were incubated with 25HOCh-coated solid-phases. Results showed maximal binding peaks at the ~ 30-50, 70-90, 110-150 and 170-190 amino acid positions (Figure 4A, black line). Similar peaks docked with minimal ΔG to 25HOCh (Figure 4A, blue line). Of the 25HOCh binding/docking peaks identified, only the 30-50 was in a similar region than the 35-47 peptide previously identified in hCRP as the main Ch-binding domain [16]. To locate the predicted interaction of 25HOCh with the CRP1-7 tridimensional structures we used PyMol. The 25HOCh docked at the CRP5 interface side with ΔG between −7.5 to −8.4 Kcal/mol (some of the contact positions at T41, E48, R71, F84, F85, S117) (Figure 4A, CRP5). In contrast, the 25HOCh docked at other CRP1-7 effector faces under the α-helix with ΔG between −8.6 to −9.1 Kcal/mol (some of the contact positions for CRP1 at R113, S115, G153, E154, Y161, and E206) (Figure 4B, CRP1). Similar ΔG values and docking location were predicted for m- or t-CRP5 (not shown). Similar docking locations were predicted for 25HOCh and Ch for most CRP1-7 within ± ΔG > ~0.5 Kcal /mol (not shown).

**Figure 4.**
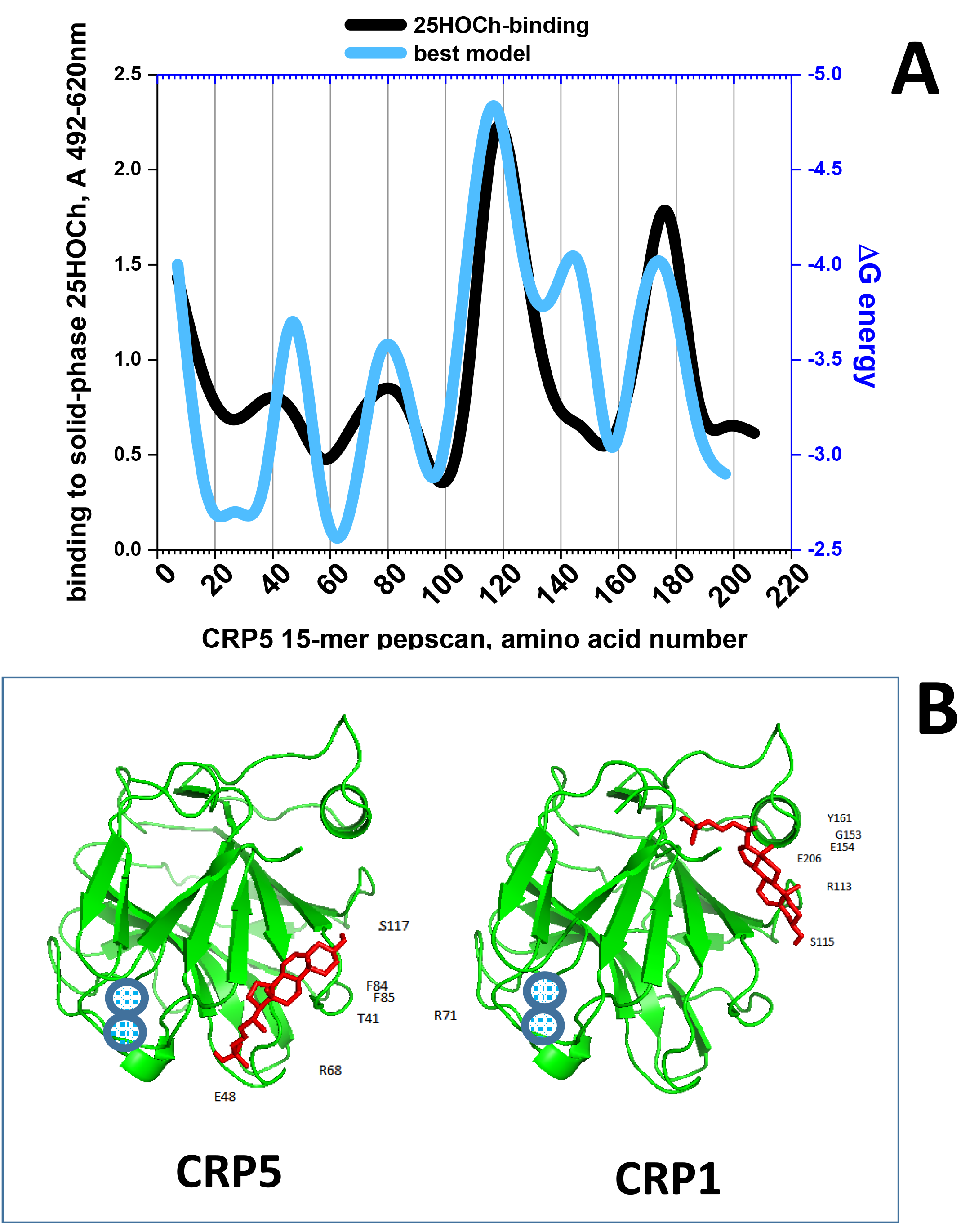
Solid-phase binding and docking predictions to 25HOCh of CRP5 pepscan peptides (A) and predicted best docking location (B) **A)** For the peptide-binding assays, a series of 15-mer peptides overlapping 5 amino acids from CRP5 were chemically synthesized by adding an amino-terminal biotin molecule. Solid-phases were coated with 2 μg per well of 25HOCh into polystyrene 96-well plates. Binding of 0.05 μg biotinylated pepscan peptides, detection with peroxidase-labelled streptavidin and staining with OPD were then performed. Means from 2 independent experiments were represented, standard deviations omitted for clarity. For the *in silico* docking predictions, the modelled pepscan peptides with the lowest energies were docked to several possible conformations of 25HOCh as described in methods. The docking energies which best fitted the binding data were then represented. **Black line**, peptide binding to 25HOCh. **Blue line**, predicted ΔG energy of peptide docking to 25HOCh. **B)** PyMOL representation of the lowest energy structures of CRP5 and CRP1 complexed to 25HOCh (the rest of CRP1-7 were similar). **Green**, CRP amino acid chains. **Red**, 25HOCh. **Blue circles**, Ca^++^ atoms located at the PC-binding pocket [6].

Therefore, both the pepscan binding and docking predictions, confirmed the existence of an interaction between 25HOCh and CRP5 and most probably for all CRP1-7.

### *In vitro* anti-SVCV effects caused by CRP1-7 in the presence of 25HOCh

Hydroxylated Chs (HOChs) are Ch oxidized derivatives with diverse biological activities, most of them correlating with inflammatory responses [49]. Among the HOChs, 25HOCh had minimal ΔG docking predictions for CRP1-7 (−8 to −9 Kcal/mol, corresponding to concentrations between 1.35 to 0.35 μM) (Figure 3). Among their biological activities, 25HOCh have been related to viral infections [24, 50], including the reduction of Spring Viremia Carp Virus (SVCV) infection in zebrafish [47]. Because of the Ch-dependence of SVCV infection (Figure 1B), the reduction of SVCV infection by zebrafish ssCRP1-7 [7] was chosen as an example of possible CRP1-7- HOChs interactions affecting the same function.

Because both 25HOCh [47] and CRP1-7 [7] have independent anti-SVCV effects, their concentrations were first titrated at different multiplicities of SVCV infection (m.o.i.) to maximize the limits of detection when they were used together. Under optimal conditions, the extent of SVCV infections (ssCRP1-7 + 25HOCh / ssCRP1-7) were further reduced 1.5 to 3-fold compared to 25HOCh (GFP+25HOCh) depending on the CRP1-7 isoform (Figure 5A). Similar results were obtained with rCRP5 and rCRP7 but not with rCRP2 (not shown). All the above commented results suggested that 25HOCh in the presence of CRP1-7 further enhanced the anti-viral effects caused by either 25HOCh or CRP1-7 alone. It is still too early to know about the mechanisms implicated, since interaction of 25HOCh with the L polymerase of SVCV [47], or 25HOCh inhibition of glycosylation in other rhabdoviruses [51], may suggest that binding to some viral proteins cannot be excluded. Furthermore, other possible interactions between 25HOCh, CRP (direct effect) and/or other CRP-induced molecules (indirect effects) present in the ssCRP1-7, may still explain the above mentioned anti-viral enhancements.

**Figure 5.**
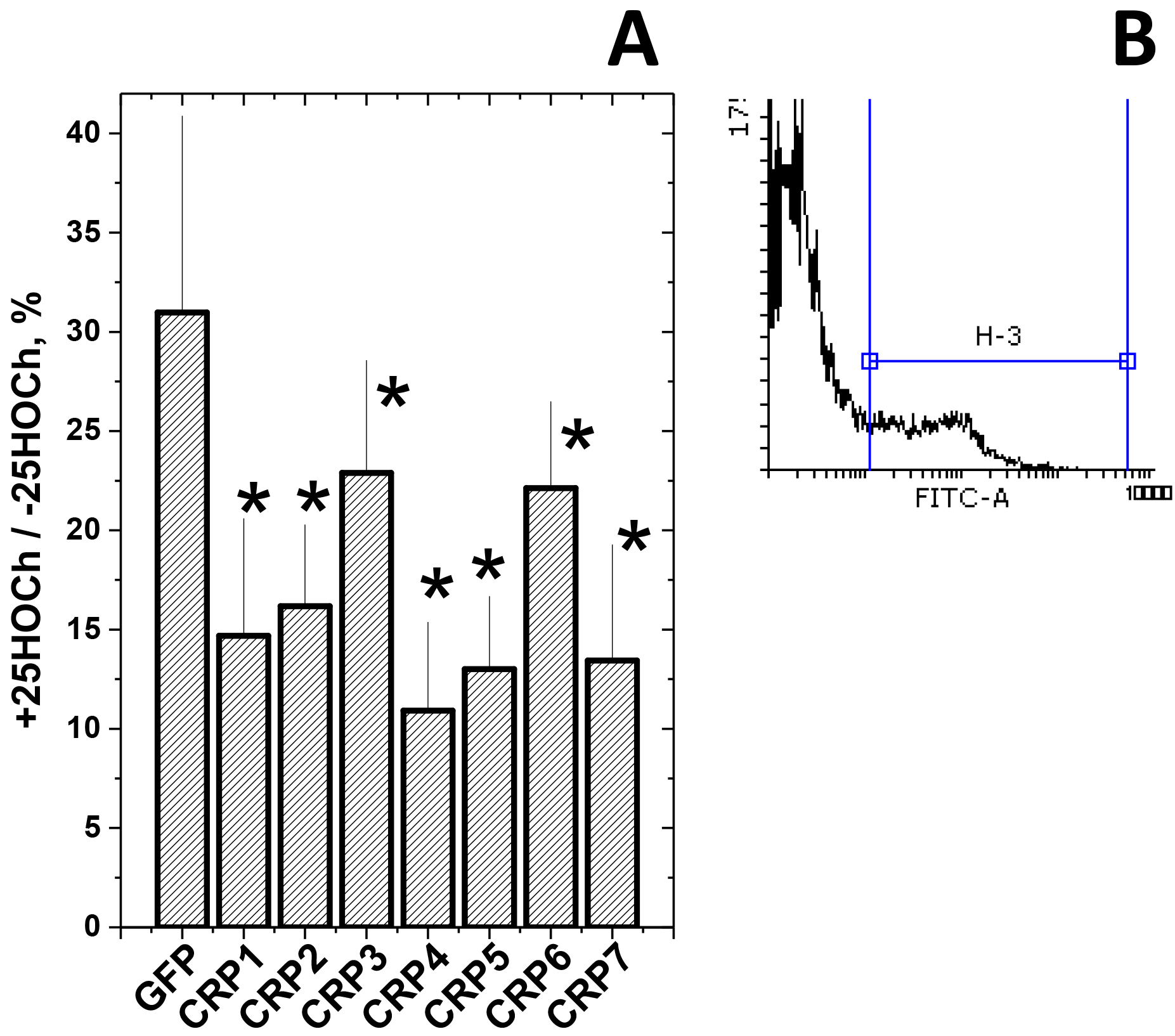
Anti-SVCV infectivity after treatment of EPC cell monolayers with 25HOCh and CRP1-7. **A)** EPC cell monolayers were incubated with 100 μl of ssGFP or ssCRP1-7 4-fold diluted in RPMI with 2% FBS ± 10 μM 25HOCh for 20 h at 26 °C. After washing, 10-2 m.o.i of SVCV were added and incubated for 24 h. After staining with anti-SVCV and fluorescein-labelled goat anti-mouse immunoglobulins [7], the number of fluorescent cells were estimated by flow cytometry. **B)** representative aspect of histograms from non-fluorescent and fluorescent cells. The number of SVCV infected EPC cells varied from 12.7 to 50.6 % (n=5), depending on the experiment. The results were expressed as relative infection percentages calculated by the formula, 100 x (number of infected cells +25HOCh / number of infected cells −25HOCh). Means and standard deviations of a representative experiment were represented (n=3). *, statistically < than cells transfected with ssGFP at p < 0.05 (Student t-test).

To explore for any possible correlations among CRP1-7 structures and 25HOCh binding or anti-viral effects, we then studied whether different oligomeric forms were present in the ssCRP1-7 employed for the above mentioned binding and anti-viral experiments.

### Insect-made rCRPs suggested different oligomerization states

*E.coli*-made zebrafish c-reactive protein CRP5 isoform (rCRP5) crystalized as trimers (t-CRP5) as shown by X-rays [6]. However, it is not yet known whether or not trimers are the physiological form or that is similar for the rest of CRP1-7 isoforms.

Our first attempts to characterize CRP1-7 isoforms included expression in *E.coli.* However, numerous experiments met with irreproducibility, expression failure, high CRP denaturation or low yields, despite reduction of autoinduction or temperature, and/or re-cloning of the best producing clones (not shown). Most probably some of those results could be explained by the toxicity of the rCRPs to *E.coli*.

Alternatively, we explored production of rCRP1 / rCRP2 / rCRP5 / CRP7 in insect cells. Results showed that while insect-made rCRP2 / rCRP5 / rCRP7 could be expressed and purified by non-denaturing affinity chromatography, all attempts to purify rCRP1 were unsuccessful. Western blot analysis using anti-polyH antibodies indicated that although small amounts of rCRP1 were present, they were not retained by the affinity columns, most probably due to polyH tail inaccessibility (not shown), perhaps becuase a different conformation of rCRP1 compared to the other rCRPs.

Polyacrylamide gel electrophoresis (PAGE) in the absence of SDS in the buffers, treating the samples under non-denaturing conditions (no heat, no SDS, no ß-mercaptoethanol and 1 mM CaCl_2_), and Western blotting with anti-p3 antibodies, showed that rCRP2 (calculated isoelectric point IP of 6.35) banded at an apparent M.W. of > 100 KDa, while rCRP5 (IP 4.6) and rCRP7 (IP 4.6) banded at ~ 75 KDa (Figure 1A left). A brief (2 min) treatment of the rCRP samples under denaturing conditions, increased the migration of all rCRP, specially that of rCRP7 (Figure 1A right). Although in the absence of SDS, the estimation of molecular weights is not accurate, the results may suggest larger sizes for rCRP2 / rCRP5 compared to rCRP7, according to previous electrophoretic data described for p-hCRP and m-hCRP [52].

In contrast, by applying PAGE in the presence of SDS in the buffers, samples under non-denaturing conditions and Western blotting, all the rCRP displayed similar bands which could be interpreted as residual amounts of trimers (~75KDa), dimers (~50 KDa) and monomers (~25 KDa) (Figure 6B, left). The amount of monomers increased when the samples were briefly treated under denaturing conditions, specially in rCRP7 were only the monomers were apparent (Figure 6B, middle). All rCRP became homogeneously monomeric (~25 KDa) when the samples were treated longer (5 min) under denaturing conditions (Figure 6B, right). The slightly different positions of the monomeric forms could be most probably due to differences in their glycosylation, although recently different post-transcriptional deimidation has been also described in cod CRPs [53].

**Figure 6.**
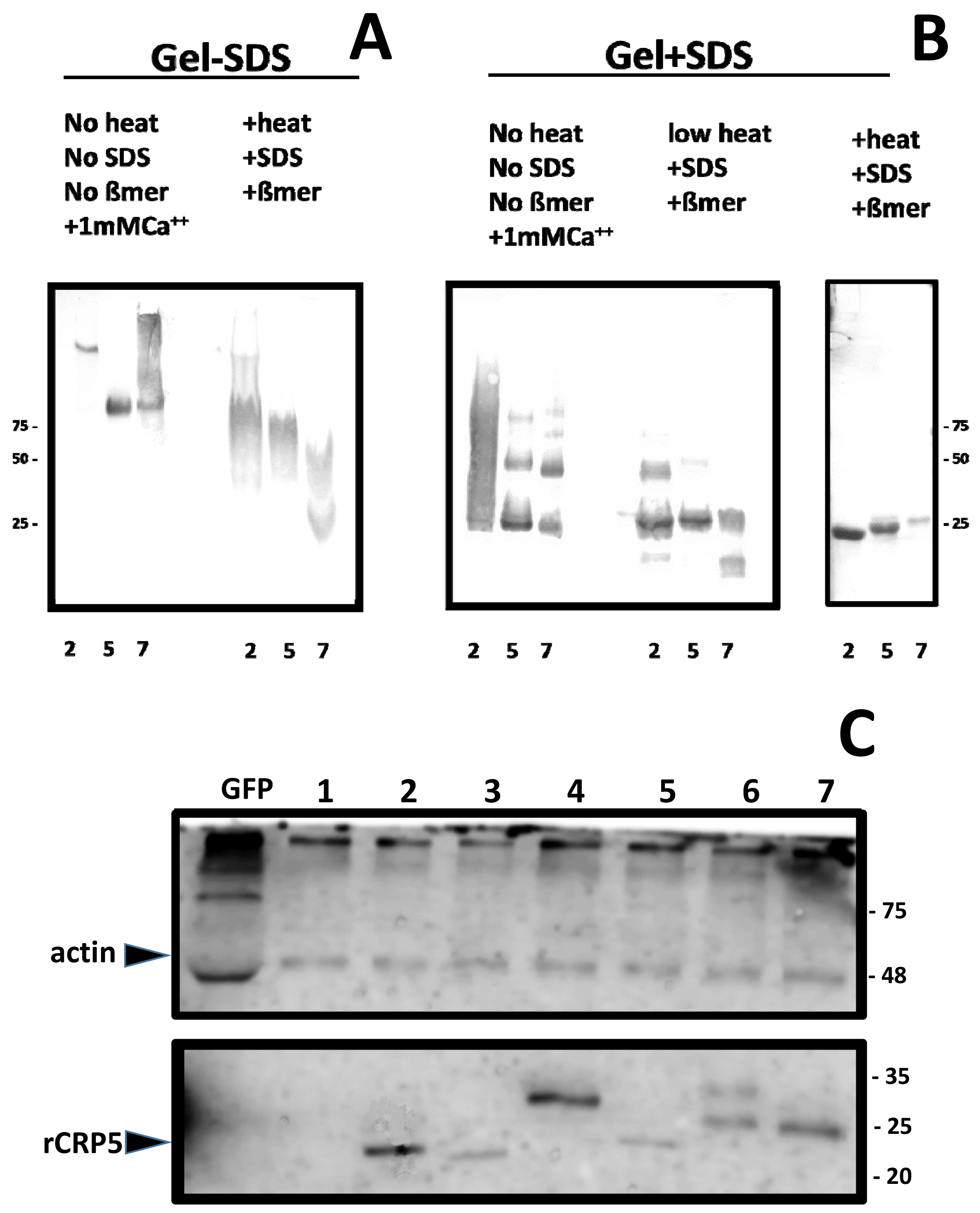
Polyacrylamide gel electrophoresis and Western blotting of rCRPs (A,B,C) and ssCRP1-7 (D). The insect-made affinity purified samples were electrophoresed in 4-20% gradient of polyacrylamide gels. **A)** Samples of rCRPs prepared and electrophoresed in the absence of SDS in the buffers and stained with Coomassie (non-denaturing conditions). **B)** Samples of rCRPs heated at 100 °C in the presence of ß-mercaptoethanol and SDS, electrophoresed in the presence of SDS in the buffers (denaturing conditions) and stained with Coomassie. **C)** Western blotting of the gel B transferred to nitrocellulose membranes, stained with anti-p3 antibody, horseradish peroxidase labelled anti-rabbit and over-exposed to diaminobenzidine (DAB) [34]. The ssCRP1-7 were electrophoresed in 15 % polyacrylamide gels. **D)** Samples of ssCRP1-7 treated at 100 °C in the presence of ß-mercaptoethanol and SDS, electrophoresed in the presence of SDS in the buffers, transferred to nitrocellulose membranes, stained with anti-actin (up) or anti-p3 (down) antibody, horseradish peroxidase labelled anti-rabbit and developed by chemiluminiscence [7]. Similar results were obtained with samples electrophoresed under non-denaturing conditions (not shown). **Numbers around the gels,** molecular weight markers in KDa. **Up left arrow**, position recognized by anti-actin antibodies. **Down left arrow**, position of purified rCRP5 recognized by anti-p3 antibodies. Results are representative of at least 3 experiments.

The most probable explanation for all the above commented data suggests that while insect-made rCRP2/rCRP5 may exist as an equilibrium among trimers, dimers and monomers, rCRP7 has a stronger tendency to form monomers.

### CRP1-7-enriched supernatants from EPC cell transfected cells appeared monomeric

Western blotting of ssCRP1-7 using anti-p3 antibodies, only detected CRP2-7 monomers of ~ 25 KDa with slightly different positions for each ssCRP2-7, with similar profiles under denaturing (Figure 6C, down),20-fold lower SDS concentration (not shown)[54] and non-denaturing (not shown) sample and buffer conditions. Similar CRP2-7 levels were present in ssCRP2-7 as shown using actin as an internal marker (Figure 6C, up). In these experiments, it was not possible to detect the presence of any CRP1 band, most probably because its lower concentration, since previous results demonstrated its presence by dot-blot when using concentrated ssCRP1 [7]. Most probably, all ssCRP1-7 were secreted from EPC transfected cells mainly as monomers.

### *In silico* predictions of zebrafish CRP1-7 tridimensional structures

To obtain more data on the possible structures of CRP1-7, their amino acid sequences were modelled using the SWISS-MODEL web program. Automatic modelling showed that only CRP2 / CRP5 rendered trimers, while the rest of the CRP1-7 modelled as monomers only (Table 2). CRP2 / CRP5 have differences in most of the modelling parameters, specially in their torsion-angle potentials, compared to other CRP1-7 (Table 2). Because the existence of ~ 70 EST from zebrafish in the UniGene Bank classified as CRP5 transcript variants [6] offered another opportunity to test the reliability of the trimer/monomer predictions mentioned above, we explored also these data. The corresponding results predicted that 97.8 % of the 47 CRP5 longest variant sequences also modelled as trimers. The comparison of the CRP5 variant amino acid sequences demonstrated 2-3-times more variations downstream of position 200 than in the rest of the molecule (Figure 7, red). On the other hand, most variations among the CRP1-7 isoforms were in the PC-binding pockets or in the hCRP-homologous Ch-binding domain (Figure 7, blue or green rectangles, respectively). Therefore, these results predicted the tendency of CRP2, CRP5 and CRP5 transcript variants to oligomerize as trimers and prompt for further studies about the biological significance for both CRP isoforms and variants.

**Table 2.** Parameter values of *in silico* predicted CRP1-7 oligomeric structures. The CRP1-7 amino acid sequences [2] were modelled as tridimensional structures using the SWISS-MODEL server with automatic template selection. In addition, 47 full-length CRP5 amino acid sequences were modelled from 73 zebrafish *crp5* EST variants (UniGene Dr.124528-Dr.162306) [6]. **QMEAN**, estimation of total similarity to the template, made up of 4 individual Z-score parameters (Cß, all-atom, solvation and torsion). The individual Z-scores compare the predicted tridimensional structures with template by: i) Cß atoms of three consecutive amino acids (**Cß**), ii) all-atoms (**AA**), iii) solvation burial status of the residues (**SO**) and iv) torsion angle potentials (**TO**). Low QMEAN score values indicate low similarity to the template. High QMEAN score values indicate high similarity to the template. **Bold**, highest and/or lowest score values. **Gray**, CRP2 / CRP5. The mean ± standard deviation (n=47) of the calculated score values of the CRP5 transcript variants were represented.

| isoform | Acc.number | Automatic SWISS prediction | QMEAN | Cβ | AA | SO | TO |
| --- | --- | --- | --- | --- | --- | --- | --- |
| CRP1 | XM_693995.4 | monomer | -0.55 | -1.36 | **-1.03** | -1.03 | 0.01 |
| CRP2 | BC097160 | homo-trimer | **0.77** | -1.64 | **-1.08** | **-0.92** | **1.19** |
| CRP3 | BC154042 | monomer | 0.01 | -1.15 | -1.15 | -1.03 | 0.51 |
| CRP4 | BC115188 | monomer | -1.81 | -1.65 | -1.95 | -1.33 | -1.07 |
| CRP5 | BC121777 | homo-trimer | **1.42** | **-0.71** | **-0.89** | **-0.60** | **1.62** |
| CRP5<br>47 variants | Dr.124528-Dr.162306 | 97.8%<br>homo-trimers | **1.12**<br>$\pm 0.31$ | **-0.90**<br>$\pm 0.5$ | **-0.99**<br>$\pm 0.2$ | **-0.68**<br>$\pm 0.4$ | **1.30**<br>$\pm 0.5$ |
| CRP6 | BC162745 | monomer | -0.45 | -1.20 | -1.17 | -1.34 | 0.15 |
| CRP7 | BC150371 | monomer | -0.42 | -1.04 | -1.22 | -1.28 | 0.15 |

**Figure 7.**
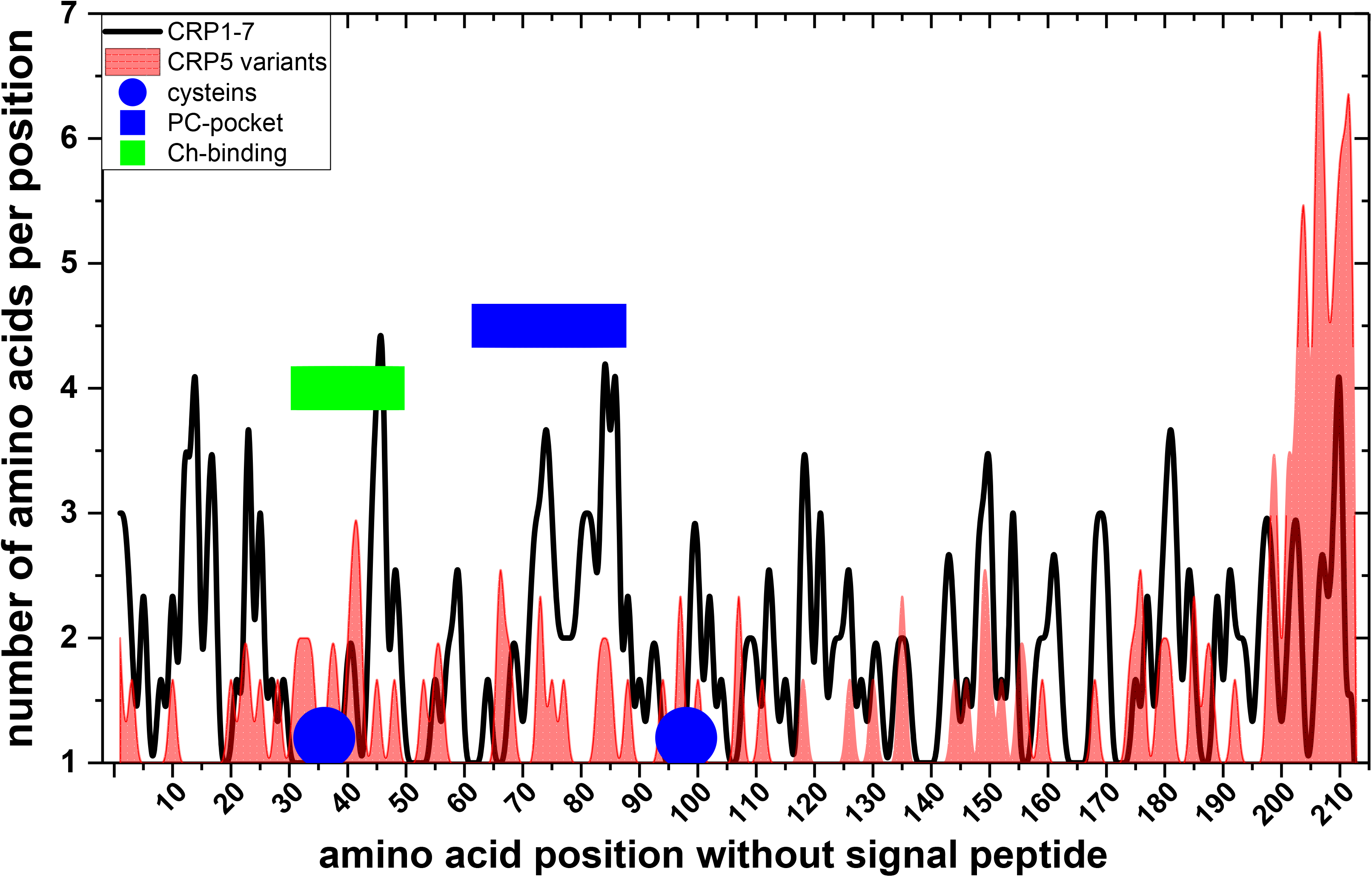
Alignement among EST-derived amino acid sequences of CRP5 transcript variants. Transcript variants corresponding to the zebrafish *crp5* gene were retrieved from 73 EST obtained from different zebrafish tissues (UniGene Dr.124528-Dr.162306) [6]. The corresponding ORFs > 100 amino acids translated by the Virtual Ribosome software (http://www.cbs.dtu.dk/services/VirtualRibosome/) were numbered without signal peptide (^1^FKNL…in CRP5) and aligned to CRP5 (BC121777). The number of different amino acids per position were represented. The data were smoothed by the cubic B-spline method (Origin Pro 2017, Northampton, MA, USA). **Blue circles**, Cysteines. **Blue rectangle**, PC-binding pocket of hCRP. **Green rectangles**, Ch-binding residues of hCRP [16]. **Black line**, number of amino acids per position of CRP1-7. **Red profile**, number of amino acids per position of CRP5 transcript variants.

The PAGE/Western data and the *in silico* predictions, together with the results of 25HOCh binding (Figure 3A) and enhancement of anti-SVCV effects (Figure 5A) by ssCRP1-7, may implicate more the m-CRP1-7 rather than t-CRP1-7 in those functions. However, CRP1-7 may also physiologically exist as an equilibrium of trimers, dimers and monomers, as shown in the cases of CRP2 /CRP5 and to lower extent in CRP7. On the other hand, because m-hCRP can also be produced *in vitro*, for instance by treatments with urea, low-pH or low-salt buffers in the absence of Ca^++^ [54, 55], the m-CRP1-7 detected in this work may have been produced by the *in vitro* manipulations (i.e., purification by affinity chromatography in the absence of Ca^++^, transfection of EPC cells, etc). We may also speculate that t-CRP1-7 could preferentially exist in fish until an stimulus triggers their conversion to m-CRP1-7 or *viceversa* since circulating hCRP is pentameric (p-hCRP) [13] and converts to monomeric (m-hCRP) after interaction with exposed phosphocholine heads and Ch-enriched lipid rafts of cellular membranes in damaged tissues [16, 17, 48, 52]. The t-CRP1-7 may be functionally analogous to the circulating p-hCRP and the m-CRP1-7 could be analogous to the converted m-hCRP. Alternatively, zebrafish m-CRP1-7 may be synthesized *de novo* as monomers. We may also think on the possibility of the existence of heterologous CRP1-7 oligomers but any of those possibilities remain speculative until specific reagents could be developed.

All together, the above commented evidence shows that the oligomeric state of CRP1-7 isoforms fine tunes their lipid binding and, at least, some of their resulting biological functionalities, as suggested before based only on their amino acid derived sequences and transcript physiology [2, 6, 7].

## Acknowledgements

Thanks are due to Paula Perez Gonzalez who helped with the experimentation. Melissa Bello-Perez was financed by the Generalidad Valenciana, fellowship ACIF/2016. This work was supported by CICYT projects AGL2014-51773-C3-R and BIO2017-82851 of the Ministerio de Economía, Industria y Competitividad of Spain.

## Competing interests

The authors declare not to have any competing interests

## Authors’ contribution

MB and AF performed the SVCV infections and westerns. JA revised the docking predictions. BN and LP collaborated in the designing of the experiments and reviewed the manuscript. JC designed and analyzed experiments, coordinate the work and drafted the manuscript. All authors read and approved the manuscript.

## List of abbreviations

BSA: bovine serum albumin
CRP: C-reactive protein
Ig: immunoglobulin
KDa: kilo Daltons
MW: molecular weight
PAGE: polyacrylamide gel electrophoresis
PBS: phosphate buffered saline
pfu: plaque forming units
SDS: sodium dodecyl sulfate
SVCV: spring viremia carp virus
VHSV: Viral haemorrhagic septicemia virus
EPC: cell line

## References

1. Bottazzi B, Inforzato A, Messa M, Barbagallo M, Magrini E, Garlanda C, et al. The pentraxins PTX3 and SAP in innate immunity, regulation of inflammation and tissue remodelling. J Hepatol. 2016;64(6):1416–27. Epub 2016/02/28. doi: S0168-8278(16)00163-X [pii] 10.1016/j.jhep.2016.02.029. PubMed PMID: 26921689.

2. Falco A, Cartwright JR, Wiegertjes GF, Hoole D. Molecular characterization and expression analysis of two new C-reactive protein genes from common carp (Cyprinus carpio). Dev Comp Immunol. 2012;37(1):127–38. Epub 2011/11/15. doi: S0145-305X(11)00278-3 [pii] 10.1016/j.dci.2011.10.005. PubMed PMID: 22079493.

3. Wang J, Tang B, Liu X, Wu X, Wang H, Xu D, et al. Increased monomeric CRP levels in acute myocardial infarction: a possible new and specific biomarker for diagnosis and severity assessment of disease. Atherosclerosis. 2015;239(2):343–9. doi: 10.1016/j.atherosclerosis.2015.01.024. PubMed PMID: 25682033.

4. Braig D, Nero TL, Koch HG, Kaiser B, Wang X, Thiele JR, et al. Transitional changes in the CRP structure lead to the exposure of proinflammatory binding sites. Nature communications. 2017;8:14188. Epub 2017/01/24. doi: ncomms14188 [pii] 10.1038/ncomms14188. PubMed PMID: 28112148.

5. McFadyen JD, Kiefer J, Braig D, Loseff-Silver J, Potempa LA, Eisenhardt SU, et al. Dissociation of C-Reactive Protein Localizes and Amplifies Inflammation: Evidence for a Direct Biological Role of C-Reactive Protein and Its Conformational Changes. Front Immunol. 2018;9:1351. doi: 10.3389/fimmu.2018.01351. PubMed PMID: 29946323; PubMed Central PMCID: PMCPMC6005900.

6. Chen R, Qi J, Yuan H, Wu Y, Hu W, Xia C. Crystal structures for short-chain pentraxin from zebrafish demonstrate a cyclic trimer with new recognition and effector faces. J Struct Biol. 2015;189(3):259–68. Epub 2015/01/17. doi: S1047-8477(15)00002-7 [pii] 10.1016/j.jsb.2015.01.001. PubMed PMID: 25592778.

7. Bello-Perez M, Falco A, Medina-Gali R, Pereiro P, Encinar JA, Novoa B, et al. Neutralization of viral infectivity by zebrafish c-reactive protein isoforms. Mol Immunol. 2017;91:145–55. doi: 10.1016/j.molimm.2017.09.005. PubMed PMID: 28915434.

8. Inforzato A, Doni A, Barajon I, Leone R, Garlanda C, Bottazzi B, et al. PTX3 as a paradigm for the interaction of pentraxins with the complement system. Semin Immunol. 2013;25(1):79–85. Epub 2013/06/12. doi: S1044-5323(13)00031-6 [pii] 10.1016/j.smim.2013.05.002. PubMed PMID: 23747040.

9. Vilahur G, Badimon L. Biological actions of pentraxins. Vascul Pharmacol. 2015;73:38–44. Epub 2015/05/13. doi: S1537-1891(15)00084-1 [pii] 10.1016/j.vph.2015.05.001. PubMed PMID: 25962566.

10. Caprio V, Badimon L, Di Napoli M, Fang WH, Ferris GR, Guo B, et al. pCRP-mCRP Dissociation Mechanisms as Potential Targets for the Development of Small-Molecule Anti-Inflammatory Chemotherapeutics. Front Immunol. 2018;9:1089. doi: 10.3389/fimmu.2018.01089. PubMed PMID: 29892284; PubMed Central PMCID: PMCPMC5985513.

11. Bang R, Marnell L, Mold C, Stein MP, Clos KT, Chivington-Buck C, et al. Analysis of binding sites in human C-reactive protein for Fc{gamma}RI, Fc{gamma}RIIA, and C1q by site-directed mutagenesis. J Biol Chem. 2005;280(26):25095–102. Epub 2005/05/10. doi: M504782200 [pii] 10.1074/jbc.M504782200. PubMed PMID: 15878871.

12. Lu J, Marjon KD, Mold C, Du Clos TW, Sun PD. Pentraxins and Fc receptors. Immunol Rev. 2012;250(1):230–8. Epub 2012/10/11. doi: 10.1111/j.1600-065X.2012.01162.x. PubMed PMID: 23046133; PubMed Central PMCID: PMC3471383.

13. Wu Y, Potempa LA, El Kebir D, Filep JG. C-reactive protein and inflammation: conformational changes affect function. Biol Chem. 2015;396(11):1181–97. Epub 2015/06/04. doi: 10.1515/hsz-2015-0149/j/bchm.just-accepted/hsz-2015-0149/hsz-2015-0149.xml [pii]. PubMed PMID: 26040008.

14. Eisenhardt SU, Habersberger J, Peter K. Monomeric C-reactive protein generation on activated platelets: the missing link between inflammation and atherothrombotic risk. Trends Cardiovasc Med. 2009;19(7):232–7. doi: 10.1016/j.tcm.2010.02.002. PubMed PMID: 20382347.

15. Eisenhardt SU, Thiele JR, Bannasch H, Stark GB, Peter K. C-reactive protein: how conformational changes influence inflammatory properties. Cell Cycle. 2009;8(23):3885–92. doi: 10.4161/cc.8.23.10068. PubMed PMID: 19887916.

16. Li HY, Wang J, Meng F, Jia ZK, Su Y, Bai QF, et al. An Intrinsically Disordered Motif Mediates Diverse Actions of Monomeric C-reactive Protein. J Biol Chem. 2016;291(16):8795–804. doi: 10.1074/jbc.M115.695023. PubMed PMID: 26907682; PubMed Central PMCID: PMCPMC4861447.

17. Wang MY, Ji SR, Bai CJ, El Kebir D, Li HY, Shi JM, et al. A redox switch in C-reactive protein modulates activation of endothelial cells. FASEB J. 2011;25(9):3186–96. doi: 10.1096/fj.11-182741. PubMed PMID: 21670067.

18. Lv JM, Lu SQ, Liu ZP, Zhang J, Gao BX, Yao ZY, et al. Conformational folding and disulfide bonding drive distinct stages of protein structure formation. Sci Rep. 2018;8(1):1494. doi: 10.1038/s41598-018-20014-y. PubMed PMID: 29367639; PubMed Central PMCID: PMCPMC5784126.

19. Bello M, Falco A, Medina R, Encinar JA, Novoa B, Perez L, et al. Structure and functionalities of the human c-reactive protein compared to the zebrafish multigene family of c-reactive-like proteins. Developmental & Comparative Immunology. 2017;69:33–40.

20. Estepa A, Coll JM. Innate multigene family memories are implicated in the viral-survivor zebrafish phenotype. Plos One. 2015;10(8):e0135483. doi: doi:10.1371/journal.pone.0135483.

21. Garcia-Valtanen P, Martinez-Lopez A, Lopez-Munoz A, Bello-Perez M, Medina-Gali RM, Ortega-Villaizan MD, et al. Zebra Fish Lacking Adaptive Immunity Acquire an Antiviral Alert State Characterized by Upregulated Gene Expression of Apoptosis, Multigene Families, and Interferon-Related Genes. Front Immunol. 2017;8:121. doi: 10.3389/fimmu.2017.00121. PubMed PMID: 28243233; PubMed Central PMCID: PMCPMC5303895.

22. Widziolek M, Prajsnar TK, Tazzyman S, Stafford GP, Potempa J, Murdoch C. Zebrafish as a new model to study effects of periodontal pathogens on cardiovascular diseases. Sci Rep. 2016;6:36023. doi: 10.1038/srep36023. PubMed PMID: 27777406; PubMed Central PMCID: PMCPMC5078774.

23. Zieden B, Kaminskas A, Kristenson M, Kucinskiene Z, Vessby B, Olsson AG, et al. Increased plasma 7 beta-hydroxycholesterol concentrations in a population with a high risk for cardiovascular disease. Arterioscler Thromb Vasc Biol. 1999;19(4):967–71. PubMed PMID: 10195924.

24. Dong H, Zhou L, Ge X, Guo X, Han J, Yang H. Antiviral effect of 25-hydroxycholesterol against porcine reproductive and respiratory syndrome virus in vitro. Antivir Ther. 2018. doi: 10.3851/IMP3232. PubMed PMID: 29561734.

25. Yang J, Gustavsson AL, Haraldsson M, Karlsson G, Norberg T, Baltzer L. High-affinity recognition of the human C-reactive protein independent of phosphocholine. Org Biomol Chem. 2017;15(21):4644–54. doi: 10.1039/c7ob00684e. PubMed PMID: 28513744.

26. Trott O, Olson AJ. AutoDock Vina: improving the speed and accuracy of docking with a new scoring function, efficient optimization, and multithreading. J Comput Chem. 2010;31(2):455–61. doi: 10.1002/jcc.21334. PubMed PMID: 19499576; PubMed Central PMCID: PMCPMC3041641.

27. Dallakyan S, Olson AJ. Small-molecule library screening by docking with PyRx. Methods Mol Biol. 2015;1263:243–50. doi: 10.1007/978-1-4939-2269-7_19. PubMed PMID: 25618350.

28. Shityakov S, Forster C. In silico predictive model to determine vector-mediated transport properties for the blood-brain barrier choline transporter. Adv Appl Bioinform Chem. 2014;7:23–36. doi: 10.2147/AABC.S63749. PubMed PMID: 25214795; PubMed Central PMCID: PMCPMC4159400.

29. Fijan N, Petrinec Z, Sulimanovic D, Zwillenberg LO. Isolation of the viral causative agent from the acute form of infectious dropsy of carp. Veterinary Archives. 1971;41:125–38.

30. ICTV. Implementation of taxon-wide non-Latinized binomial species names in the family *Rhabdoviridae*. Rhabdoviridae Study Group. 2015:9.

31. Perez-Filgueira DM, Resino-Talavan P, Cubillos C, Angulo I, Barderas MG, Barcena J, et al. Development of a low-cost, insect larvae-derived recombinant subunit vaccine against RHDV. Virology. 2007;364:422–30.

32. Smith PK, Krohn RI, Hermanson GT, Mallia AK, Gartner FH, Provenzano MD, et al. Measurement of protein using bicinchoninic acid. Anal Biochem. 1985;150(1):76–85. Epub 1985/10/01. doi: 0003-2697(85)90442-7 [pii]. PubMed PMID: 3843705.

33. Biro A, Cervenak L, Balogh A, Lorincz A, Uray K, Horvath A, et al. Novel anti-cholesterol monoclonal immunoglobulin G antibodies as probes and potential modulators of membrane raft-dependent immune functions. J Lipid Res. 2007;48(1):19–29. doi: 10.1194/jlr.M600158-JLR200. PubMed PMID: 17023738.

34. Torrent F, Villena A, Lee PA, Fuchs W, Bergmann SM, Coll JM. The amino-terminal domain of ORF149 of koi herpesvirus is preferentially targeted by IgM from carp populations surviving infection. Arch Virol. 2016;161(10):2653–65. doi: 10.1007/s00705-016-2934-4. PubMed PMID: 27383208.

35. Coll JM. Herpesvirus infection induces both specific and hetrologous anti-viral antibodies in carp. Frontiers in Immunology. 2018;9. doi: doi: 10.3389/fimmu.2018.00039.

36. Neron B, Menager H, Maufrais C, Joly N, Maupetit J, Letort S, et al. Mobyle: a new full web bioinformatics framework. Bioinformatics. 2009;25(22):3005–11. doi: 10.1093/bioinformatics/btp493. PubMed PMID: 19689959; PubMed Central PMCID: PMCPMC2773253.

37. Estepa A, Coll JM. Pepscan mapping and fusion related properties of the major phosphatidylserine-binding domain of the glycoprotein of viral hemorrhagic septicemia virus, a salmonid rhabdovirus. Virology. 1996;216:60–70.

38. Estepa AM, Rocha AI, Mas V, Perez L, Encinar JA, Nunez E, et al. A protein G fragment from the Salmonid viral hemorrhagic septicemia rhabdovirus induces cell-to-cell fusion and membrane phosphatidylserine translocation at low pH. Journal of Biological Chemistry. 2001;276(49):46268–75. PubMed PMID: ISI:000172573100106.

39. Biasini M, Bienert S, Waterhouse A, Arnold K, Studer G, Schmidt T, et al. SWISS-MODEL: modelling protein tertiary and quaternary structure using evolutionary information. Nucleic Acids Res. 2014;42(Web Server issue):W252–8. doi: 10.1093/nar/gku340. PubMed PMID: 24782522; PubMed Central PMCID: PMCPMC4086089.

40. Arnold K, Bordoli L, Kopp J, Schwede T. The SWISS-MODEL workspace: a web-based environment for protein structure homology modelling. Bioinformatics. 2006;22(2):195–201. doi: 10.1093/bioinformatics/bti770. PubMed PMID: 16301204.

41. Guex N, Peitsch MC. SWISS-MODEL and the Swiss-PdbViewer: an environment for comparative protein modeling. Electrophoresis. 1997;18(15):2714–23. doi: 10.1002/elps.1150181505. PubMed PMID: 9504803.

42. Mariani V, Kiefer F, Schmidt T, Haas J, Schwede T. Assessment of template based protein structure predictions in CASP9. Proteins. 2011;79 Suppl 10:37–58. doi: 10.1002/prot.23177. PubMed PMID: 22002823.

43. Tanaka T, Robey FA. A new sensitive assay for the calcium-dependent binding of C-reactive protein to phosphorylcholine. J Immunol Methods. 1983;65(3):333–41. PubMed PMID: 6361145.

44. Pepys MB. C-reactive protein fifty years on. Lancet. 1981;1(8221):653–7. PubMed PMID: 6110874.

45. Agrawal A, Xu Y, Ansardi D, Macon KJ, Volanakis JE. Probing the phosphocholine-binding site of human C-reactive protein by site-directed mutagenesis. J Biol Chem. 1992;267(35):25353–8. Epub 1992/12/15. PubMed PMID: 1460031.

46. Yang Q, Zhang Q, Tang J, Feng WH. Lipid rafts both in cellular membrane and viral envelope are critical for PRRSV efficient infection. Virology. 2015;484:170–80. doi: 10.1016/j.virol.2015.06.005. PubMed PMID: 26115164.

47. Pereiro P, Forn-Cuni G, Dios S, Coll J, Figueras A, Novoa B. Interferon-independent antiviral activity of 25-hydroxycholesterol in a teleost fish. Antiviral Res. 2017;145:146–59. doi: 10.1016/j.antiviral.2017.08.003. PubMed PMID: 28789986.

48. Ji SR, Wu Y, Zhu L, Potempa LA, Sheng FL, Lu W, et al. Cell membranes and liposomes dissociate C-reactive protein (CRP) to form a new, biologically active structural intermediate: mCRP(m). FASEB J. 2007;21(1):284–94. doi: 10.1096/fj.06-6722com. PubMed PMID: 17116742.

49. Gold ES, Diercks AH, Podolsky I, Podyminogin RL, Askovich PS, Treuting PM, et al. 25-Hydroxycholesterol acts as an amplifier of inflammatory signaling. Proc Natl Acad Sci U S A. 2014;111(29):10666–71. doi: 10.1073/pnas.1404271111. PubMed PMID: 24994901; PubMed Central PMCID: PMCPMC4115544.

50. Civra A, Cagno V, Donalisio M, Biasi F, Leonarduzzi G, Poli G, et al. Inhibition of pathogenic non-enveloped viruses by 25-hydroxycholesterol and 27-hydroxycholesterol. Sci Rep. 2014;4:7487. doi: 10.1038/srep07487. PubMed PMID: 25501851; PubMed Central PMCID: PMCPMC4265783.

51. Shrivastava-Ranjan P, Bergeron E, Chakrabarti AK, Albarino CG, Flint M, Nichol ST, et al. 25-Hydroxycholesterol Inhibition of Lassa Virus Infection through Aberrant GP1 Glycosylation. mBio. 2016;7(6). doi: 10.1128/mBio.01808-16. PubMed PMID: 27999160; PubMed Central PMCID: PMCPMC5181775.

52. Potempa LA, Yao ZY, Ji SR, Filep JG, Wu Y. Solubilization and purification of recombinant modified C-reactive protein from inclusion bodies using reversible anhydride modification. Biophys Rep. 2015;1:18–33. doi: 10.1007/s41048-015-0003-2. PubMed PMID: 26942216; PubMed Central PMCID: PMCPMC4762138.

53. Magnadottir B, Hayes P, Gisladottir B, Bragason B, Hristova M, Nicholas AP, et al. Pentraxins CRP-I and CRP-II are post-translationally deiminated and differ in tissue specificity in cod (Gadus morhua L.) ontogeny. Dev Comp Immunol. 2018;87:1–11. doi: 10.1016/j.dci.2018.05.014. PubMed PMID: 29777721.

54. Taylor KE, van den Berg CW. Structural and functional comparison of native pentameric, denatured monomeric and biotinylated C-reactive protein. Immunology. 2007;120(3):404–11. doi: 10.1111/j.1365-2567.2006.02516.x. PubMed PMID: 17163961; PubMed Central PMCID: PMCPMC2265887.

55. Potempa LA, Maldonado BA, Laurent P, Zemel ES, Gewurz H. Antigenic, electrophoretic and binding alterations of human C-reactive protein modified selectively in the absence of calcium. Mol Immunol. 1983;20(11):1165–75. PubMed PMID: 6656768.

